# Differences in other-directed emotion regulation tracks connectivity between amygdala and prefrontal regions during fairness decisions

**DOI:** 10.64898/2026.05.14.724908

**Authors:** Melanie Kos, Yi (Jen) Yang, Chelsea Helion, David V. Smith

**Affiliations:** Department of Psychology and Neuroscience, Temple University, Philadelphia, Pennsylvania, United States

**Keywords:** emotion regulation, fairness, social decision-making

## Abstract

Fairness decisions often integrate affective responses within a social context, yet emotion regulation in this literature has been largely studied as a self-directed process rather than an interpersonal one. We examined how individual differences in other-directed emotion regulation—measured with the Emotion Regulation of Others and Self (EROS) scale—relate to behavioral and neural responses during fairness decisions in 138 adults completing a variant of the Ultimatum Game with human and computer partners during fMRI. Behaviorally, participants who more strongly endorsed worsening others’ emotions rejected unfair offers more frequently, and this tendency interacted with offer fairness to amplify rejection of unfair offers. At the neural level, the left anterior insula tracked offer unfairness more strongly in social versus nonsocial contexts, consistent with sociality modulating the neural encoding of fairness. Right dlPFC activation during socially unfair offers was greater among individuals who preferred to improve others’ emotions. Connectivity analyses revealed that social fairness sensitivity predicted stronger amygdala–orbitofrontal and amygdala–dmPFC coupling; the latter was further amplified among individuals higher in other-directed emotion worsening. Together, these findings identify interpersonal emotion regulation as an understudied source of variation in the affective and prefrontal systems supporting fairness-based social decisions.

## Introduction

Whether it is splitting the bill for a meal evenly despite having ordered the least or allowing a harried traveler to cut the airport security line, people sometimes choose to accept outcomes that feel unfair. Individuals vary in how strongly they respond to situational unfairness, or what we term sensitivity to fairness (Gu et al., 2015; Shamay-Tsoory et al., 2012). Individual differences in sensitivity to fairness may influence the intensity and type of emotions elicited by perceived unfairness. However, the negative affective state elicited by unfairness can be changed via emotion regulation processes that serve to modulate affect. To date, much of the research on emotion regulation in decision-making has focused on it as a self-directed or intrapersonal process (Buhle et al., 2014; Denny et al., 2023; Morawetz et al., 2020). However, many (if not most) emotionally impactful decisions occur in interactions with other social agents, where individuals must not only manage their own emotional responses, but also actively infer and respond to the emotional states of others (Reddan et al., 2025). Here, we examine the interplay between sensitivity to fairness, other-directed emotion regulation, and social decision-making using a variant of the Ultimatum Game (UG) completed during functional neuroimaging.

Within neuroeconomics, perceptions of fairness and decision-making have often been measured using the UG, a two-player bargaining task where one person (the proposer) proposes how to split an endowment and the other (the receiver) accepts or rejects the proposal. Although the economically rational response is to accept any positive offer, receivers routinely reject unfair offers at a cost to themselves (Fehr & Schmidt, 1999; Rǎțalǎ & Sanfey, 2026). These rejections reflect social preferences, indicating that individuals weigh others’ outcomes, intentions, and welfare alongside their own financial gain. Notably, the social context of the interaction further shapes these processes. Unfair offers from human proposers are rejected more frequently than equivalent offers from computer proposers, suggesting that interpersonal social contexts may alter affective responses to perceived unfairness and thus influence subsequent decision-making (Sanfey et al., 2003; van ’t Wout et al., 2006; Luo et al., 2014; T. Wu et al., 2013; Swiderska et al., 2019). These context-sensitive rejection patterns may be supported by brain areas associated with affect and social decision-making, such as the amygdala and anterior insula, with neural responses in these regions potentially reflecting the social preferences that ultimately drive rejection decisions.

Individual differences in emotion regulation, or modulating one’s affective state, can also be a significant contributor to decision-making processes (Schmidt & Helion, 2026). Decision-making paradigms that result in increased negative emotions, such as anger or disgust, are associated with further increases in rejection behavior to unfair offers as compared to baseline (Ma et al., 2012; Gan et al., 2025). Moreover, in a UG paradigm, instructed down-regulation of negative affect increases acceptance of unfair offers and can promote more prosocial responses toward previously unfair partners (Grecucci et al., 2013). Taken together, this suggests that unchecked negative emotion, and its self-directed regulation, both impact social decision-making. What remains unclear is whether tendencies to influence *others’* emotions may similarly influence social decision-making when one perceives and/or is sensitive to unfair treatment.

Prior work in the social decision-making space has identified key limbic and cortical regions associated with social decision-making, emotion generation, and emotion regulation (Q. Wu & O’Doherty, 2026). In the UG, increased activation in both the amygdala and anterior insula has been associated with heightened negative affective responses and greater rejection of inequitable offers (Sanfey et al., 2003; Tabibnia et al., 2008; Harlé & Sanfey, 2012; Cheng et al., 2017). The overlapping contributions of these two regions in supporting social preferences, fairness evaluation, and emotion regulation raises the question of whether individual differences in other-directed emotion regulation tendencies reflect how these systems are recruited and coordinate with other regions during social decision-making. Indeed, both the amygdala and anterior insula are noted to have efferent connections to orbitofrontal and medial prefrontal cortices (Molnar-Szakacs & Uddin, 2022; Kim et al., 2011; Sun et al., 2023) – regions that are noted for their involvement in social cognition and emotion regulation processes (Amodio & Frith, 2006; Gilead et al., 2016) and have been associated with rejection of unfair offers in the UG (Cheng et al., 2017; Feng et al., 2015; Gan et al., 2025; Grecucci et al., 2013; Speitel et al., 2019).

The present study aims to identify how individual differences in sensitivity to fairness and other-directed emotion regulation interact in the context of social and non-social decision-making. To do so, we collected behavioral and functional neuroimaging data during a one-shot Ultimatum Game with both human and computer partners. We predicted that individuals with a greater tendency to worsen others’ emotions would show heightened behavioral sensitivity to fairness, especially in social vs. nonsocial contexts, reflected as an increased rejection of unfair offers from a human as compared to the computer. At the neural level, we expected that differences in partner condition and fairness would increasingly recruit regions such as the amygdala and anterior insula, consistent with the notion that these regions encode information about social context as well as social preferences. Finally, we predicted that individual differences in behavioral sensitivity to fairness and/or other-directed emotion regulation tendencies will be associated with or moderate activation and connectivity in neural regions implicated in affective and social evaluation.

## Methods

### Participants

We recruited 225 adults from the greater Philadelphia metro area (ages 21-84) as part of a larger project focused on decision-making across the lifespan (Figure 1, (D. V. Smith et al., 2024). Prescreening excluded individuals with magnetic resonance (MR) contraindications, psychiatric or neurological disorders, or medical conditions that would preclude their ability to complete the study. To reduce potential confounds associated with age-related cognitive changes, sixty-one individuals aged 56 or older were excluded from all analyses. An additional 26 participants were excluded due to insufficient or missing task, questionnaire, or imaging data. Participants provided informed consent in a manner approved by Temple University’s Institutional Review Board and were compensated for their participation. The final sample used for analyses included 138 participants aged 21 to 55 (88 female; 45 male; 5 non-binary; mean age = 33.7 years, SD = 9.73). Demographic information is reported in Supplemental Table 1.

**Figure 1.**
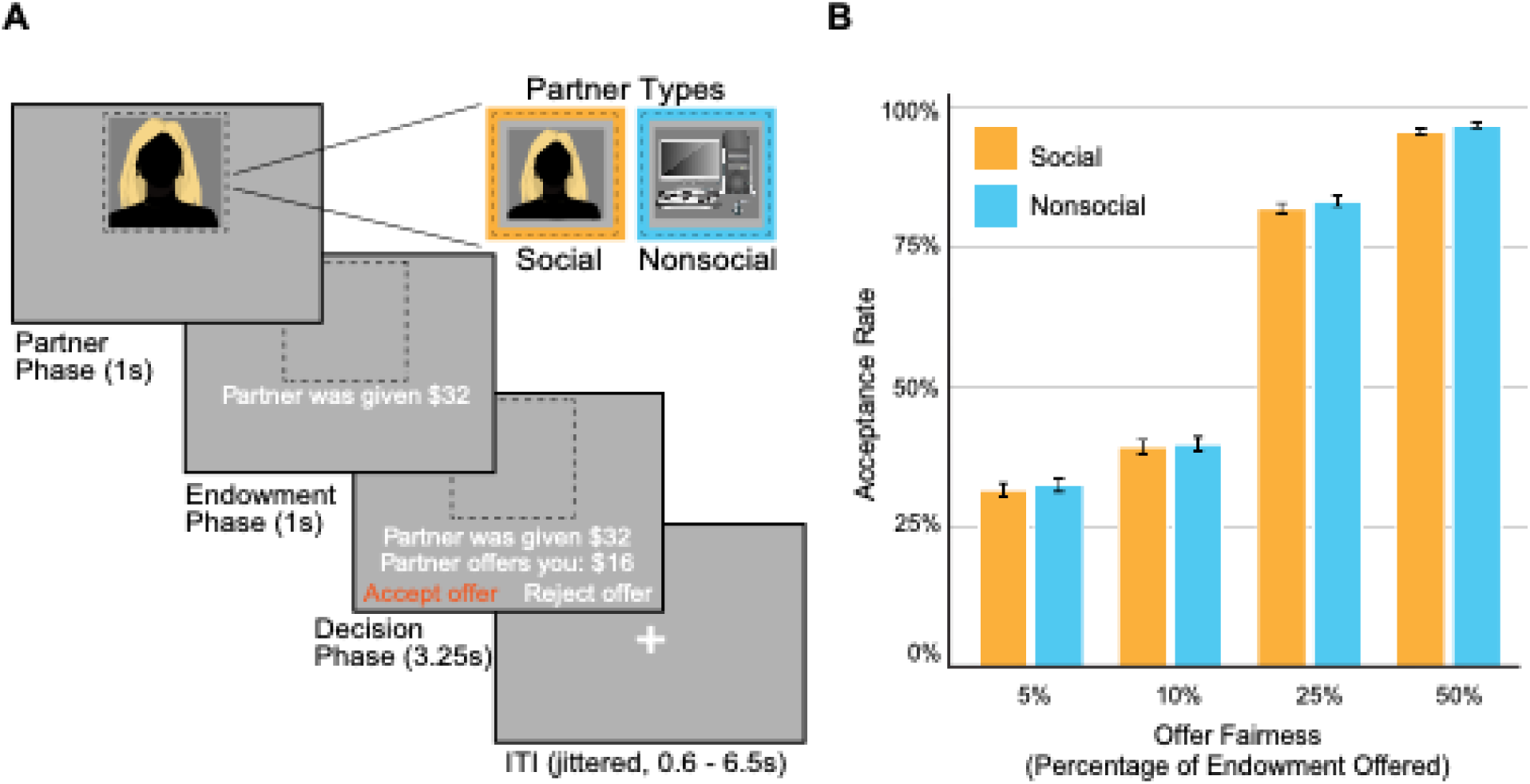
Scanner task and behavioral results. Each trial of the Ultimatum Game consisted of three sequential phases: 1) presentation of the proposer partner’s photo, 2) presentation of the monetary endowment given to the proposer, and 3) offer decision, during which participants saw the proposed split (i.e., 5-50% of the endowment) and accepted or rejected via button box response. The partner and endowment phase were presented for 1 second each, followed by the 3.25 second offer decision phase. Intertrial intervals were jittered to be between 0.6 and 6.5 seconds. Panel B, demonstrates participants’ behavioral acceptance rates across the four offer categories, where fair offers were accepted significantly more frequently than unfair offers, replicating the behavioral sensitivity to fairness effect. Acceptance rates did not significantly differ based on the offer originating from the social (human stranger) or nonsocial (computer) partner.

### Tasks

All participants completed the Ultimatum Game (UG) as the receiver while undergoing MR scanning. The UG task consisted of two runs of 48 one-shot trials each. Scanning runs where subjects missed 25% or more of trials were excluded from analyses. Participants played in two potential partner conditions: social and nonsocial. In the social condition, participants were informed that their partner was a stranger (i.e., a past study participant). In the nonsocial condition, they were told that partner was a computer (i.e., an algorithm generating procedural responses). Additionally, each trial had one of two possible endowment amounts: $16 (low) or $32 (high). During each trial, participants were sequentially shown an image of their partner, the endowment amount their partner received, followed by the amount their partner decided to offer to share (Figure 1). Participants then decided to accept or reject this offer, knowing that if they rejected the offer, neither individual would receive any money. Participants were told that one of the trials would be selected at random and added to their bonus payment. They were also informed that proposer partners would receive payment based on that random trial’s outcome, with any money won either going directly to the social partner or being returned to a pool of laboratory funds for outcomes with nonsocial partners.

### Individual Difference Measures

#### Emotion Regulation

Prior to MR scanning, participants completed a battery of self-report measures including basic demographics (e.g., age, gender, race) and emotion regulation tendency. Emotion regulation tendencies were assessed using the 19-question version of the Emotion Regulation of Others and Self Scale (Niven et al., 2011), which measures how individuals try to regulate their own and others’ emotions, either to improve or worsen affect. Items were rated on a 5-point Likert scale from 1 (Not at all) to 5 (A great deal). Responses were used to compute four subscale scores reflecting individuals’ average tendencies to 1) improve one’s own emotions (*Intrinsic Affect-Improving*), 2) worsen one’s own emotions (*Intrinsic Affect-Worsening*), 3) improve others’ emotions (*Extrinsic Affect-Improving,* or other-directed emotion improving), and 4) worsen others’ emotions (*Extrinsic Affect-Worsening*, or other-directed emotion worsening). Scores were then averaged across responses on each subscale and grand-mean centered prior to inclusion in models.

#### Sensitivity to Fairness

To examine how decisions to accept or reject offers that vary in fairness and agent sociality may be associated with emotion regulation abilities and/or neural activation, we first computed individuals’ behavioral sensitivity to fairness. Offers were binned into four fairness levels, reflecting a percentage of the total offer allotted to the responder: 5%, 10%, 25%, and 50%. To ensure offer amounts were in whole dollar amounts, some percentage levels were slightly adjusted (see Supplemental). Per-participant slopes for behavioral sensitivity to fairness (estimated separately for social, nonsocial, and combined trials) were derived from a generalized linear mixed effects model with trial-level fairness as a continuous predictor and a random intercept for participant; full specification in the Supplemental. A composite difference score was also computed for each participant by subtracting their nonsocial slope from their social slope, reflecting their sensitivity to fairness in social versus nonsocial contexts (See Supplemental for more details). These slopes reflect the change in the log-odds of accepting an offer per-unit increase in fairness on a 0-1 scale. For example, a higher value would reflect an increased sensitivity to fairness, and thus a higher likelihood of rejecting unfair offers and accepting fair ones. All-trial and social vs. nonsocial extracted slope values served as dependent variables in our behavioral analyses and as predictor variables in our neural analyses. All GLMMs were estimated using the glmer function from the “lme4” package in R (Bates et al., 2015).

### Neuroimaging Data Acquisition and Preprocessing

All images were collected using a 3T Siemens PRISMA MRI scanner (Siemens AG, Muenchen, Germany) with a 20-channel head coil at the Temple University Brain Research and Imaging Center. Two scanning runs of 240 volumes each were collected per participant. Functional MRI data were acquired using a gradient echo-planar imaging (EPI) with four echoes (13.8, 31.54, 49.28, 67.02 ms) and simultaneous multislice (multiband factor = 3) with in-plane acceleration (GRAPPA = 2) and partial Fourier set to 7/8. This multi-echo sequence was collected with the following parameters: matrix = 80×80; 2.7 mm^3^ isotropic resolution; 10% gap between slices; 51 axial slices; TR = 1615 ms (D. V. Smith et al., 2024).

Neuroimaging data were converted to the Brain Imaging Data Structure (BIDS) format using HeuDiConv (Halchenko et al., 2024), defaced used PyDeface (Gulban et al., 2022), field maps and distortion correction applied via warpkit (Van et al., 2023), and then minimally processed using the default pipeline provided in fMRIPrep 24.1.1 (Esteban et al., 2019; Markiewicz et al., 2024). Further, spatial smoothing was performed using a 5mm FHWM Gaussian smoothing kernel with FMRI Expert Analysis Tool (FEAT) from the FMRIB Software Library (FSL) (Woolrich et al., 2001). MRIQC was conducted with the full dataset of 225 subjects, which flagged 19 subject runs for exclusion due to poor neuroimaging data quality (Esteban 2017). Additionally, TE-dependence analysis was performed on input data to obtain additional confounds using the TEDANA workflow (DuPre et al., 2021). Full preprocessing details can be found in the Supplemental.

### Neuroimaging Analyses

Neuroimaging analyses were conducted using FSL version 6.0.7.16 (S. M. Smith et al., 2004; Jenkinson et al., 2012). First-level models were estimated using FSL’s FEAT framework with FILM prewhitening to account for local autocorrelation (Woolrich et al., 2001). We constructed two types of models: 1) an activation model to estimate BOLD responses during the decision phase of the Ultimatum Game and 2) a psychophysiological interaction (PPI) model to examine task-dependent changes in connectivity during the decision phase. The activation model included 11 task-related regressors: four regressors reflecting each combination of endowment size (high vs. low) and partner type (stranger vs. computer), four corresponding regressors parametrically modulated by offer fairness, two regressors for response time (one constant, one parametrically modulated), and one regressor for missed trials. All task-related regressors were convolved with FSL’s canonical hemodynamic response function.

PPI seeds for the bilateral amygdala and left anterior insula (AIns) were identified from activation-based analyses. The PPI models included the same 11 task regressors described above, a regressor for the mean time course within the PPI seed, and 11 interaction terms reflecting the product of the seed time course and each task regressor, yielding 23 total regressors. These models allowed us to examine how seed connectivity during decision-making varied as a function of social context, offer fairness, and individual differences in emotion regulation.

Both the activation and connectivity models included nuisance regressors to account for motion and physiological noise. These nuisance regressors consisted of six motion parameters (translations and rotations), the first six aCompCor components explaining the most variance, two non-steady state volumes, and framewise displacement (FD). High-pass filtering was applied using a 128-second cut-off with discrete cosine basis functions. Additional nuisance regressors were included to account for non-BOLD-related variance, including head motion artifacts and components identified by TEDANA (the tedana Community et al., 2024).

Within each participant, functional data from both runs were combined using fixed-effects modeling. Across participants, data were analyzed using mixed-effects models in FLAME Stage 1 with outlier de-weighting (Woolrich et al., 2004). Group-level analyses focused on whether neural responses to fairness differed by partner type (i.e., sensitivity to fairness for the social vs. nonsocial condition), and whether these effects were moderated by emotion regulation tendencies. One model tested the effect of sensitivity to fairness and four additional models tested the interaction between sensitivity to fairness and each EROS subscale. All models included age, gender, framewise displacement (FD), and temporal signal-to-noise ratio (tSNR) as covariates. Z-statistic images were thresholded using a cluster-forming threshold of Z > 3.1 and corrected at the cluster level using Gaussian Random Field Theory (Woo et al., 2014), with a family-wise error rate of p < .0125 to account for the four EROS-related comparisons. All neuroimaging results were visualized using MRIcroGL (Rorden, 2025).

## Results

### Accept behavior is predicted by offer fairness and moderated by extrinsic affect-worsening

We first examined the effects of offer fairness and partner identity (i.e., social vs. non-social) on offer acceptance rates using multilevel logistic regression. The analyses revealed a significant positive main effect of offer fairness (*b* = 15.80, *SE* = 0.34, *z* = 47.0, p < .001; Supplemental Figure 1), indicating that participants were more likely to accept an offer as its fairness increased. We found no significant interaction between offer fairness and partner identity, nor a main effect of partner identity (i.e., social vs. non-social). These results suggest that fairness strongly influenced acceptance behavior, and this effect was not modulated by whether the partner was a social or nonsocial agent. We next tested whether emotion regulation tendencies moderated this relationship between offer fairness and participants’ acceptance behavior; at the trial-level, extrinsic affect-worsening, extrinsic affect-improving, and intrinsic affect-worsening were significant (Supplemental Figure 2).

To test whether individual differences in emotion regulation abilities moderated responses elicited by individual differences in sensitivity to fairness, we ran four GLMMs with a random intercept for subject and a random slope for offer fairness. Each model included an interaction term between offer fairness and one of the four EROS subscales. Only the extrinsic affect-worsening model demonstrated a significant interaction with offer fairness, though this does not pass Bonferroni nor Benjamini-Hochberg multiple comparison correction (*b* = 5.25, *SE* = 2.40, *z* = 2.19, *p* = .029; Supplemental Figure 3). Individuals who reported making greater efforts to worsen others’ moods exhibited higher behavioral sensitivity to fairness by being more likely to reject unfair offers. None of the other emotion regulation dimensions significantly moderated behavioral sensitivity to fairness. Next, we sought to examine whether these individual differences could explain acceptance differences in responses based on partner condition but found no significant differences across all models. Finally, we investigated whether these predictors moderated participants’ responses to offer fairness across social and nonsocial partners but found no significant three-way interactions. Together, these results replicate previous findings that fairness strongly predicts acceptance behavior and adds that individuals’ other-focused emotion worsening tendencies reliably amplify this fairness effect.

### Fairness-related activation in right dlPFC varies with tendencies to improve others’ emotions

We first compared trials involving social partners to those involving nonsocial partners (social > nonsocial). In this whole-brain contrast, we observed greater activation in regions associated with social cognition, including the bilateral amygdala, fusiform face area, medial prefrontal cortex, and temporal parietal junction (Supplemental Table 2). We then tested whether neural responses tracked fairness by using a whole-brain contrast of parametrically modulated offer fairness, where the left anterior insula was found to vary as a function of offer fairness (Supplemental Table 3). To quantify this effect, we extracted left anterior insula activation across the categorical levels of offer fairness (e.g., 5-50%) and across social and nonsocial contexts. A linear mixed-effects model revealed significant main effects of fairness level, *F*(3, 959) = 14.95, *p* < .001, and partner condition, *F*(1, 959) = 33.22, *p* < .001, as well as a significant interaction, *F*(3, 959) = 2.81, *p* = .028. These results indicate that the anterior insula tracks both offer unfairness and social context, and anterior insula sensitivity to fairness is modulated by partner sociality (Figure 2).

**Figure 2.**
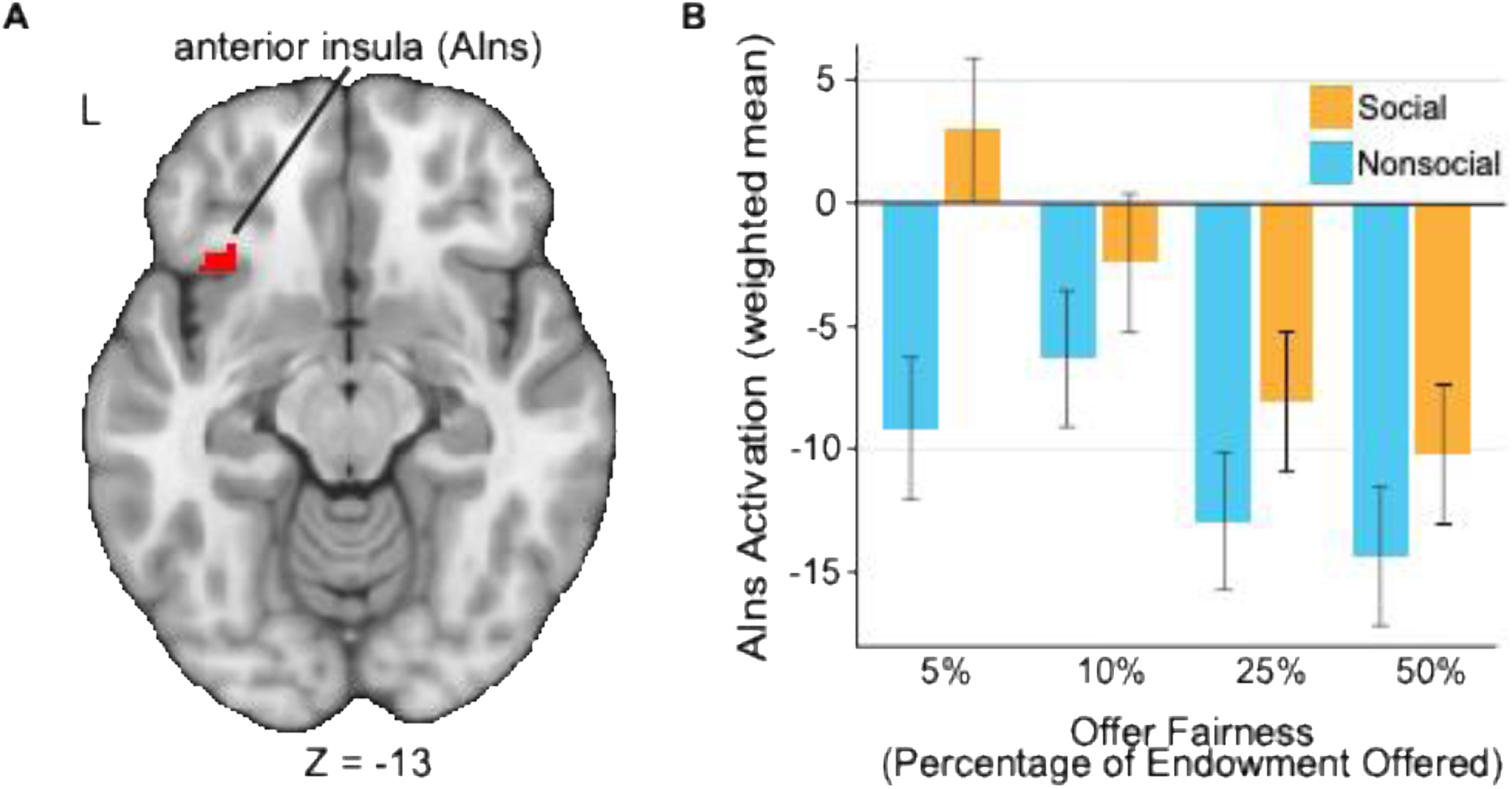
Offer fairness-related anterior insula activation in social vs. nonsocial contexts. Panel A displays the left anterior insula (AIns) region identified in a parametrically modulated fairness-related voxel-wise thresholded activation contrast where unfair offers evoked differential neural activation (*k* = 13, MNI = -32, 23, -13, *z* = 6.15; Mask: https://identifiers.org/neurovault.image:1023930 and unthresholded results used to derive mask: https://identifiers.org/neurovault.image:1028055). Panel B shows the extracted activation values across the four fairness levels for the social and nonsocial partner conditions, demonstrating context-dependent neural sensitivity to fairness.

Given that brain activation varied with both social context and fairness, we next examined whether individual differences in participants’ emotion regulation tendencies altered the strength of these effects. Whole-brain activation analyses identified a significant moderation effect of extrinsic affect-improving in the right dorsolateral prefrontal cortex (dlPFC, *k* = 38, *z* = 4.4, *p* < .002, adjusted *p* = .005; Figure 3). This indicates that participants who were more sensitive to fairness in the social condition and more inclined to improve others’ mood demonstrated increased dlPFC activation relative to those lower in other-directed emotion improving. There were no significant effects observed for any of the other three emotion regulation tendencies assessed. These results demonstrate that while neural responses to social context and fairness were largely consistent across participants, individual differences in other-directed emotion improving tendencies were associated with increased dlPFC sensitivity to fairness in social contexts, linking interpersonal emotion regulation tendencies to the enforcement of social norms.

**Figure 3.**
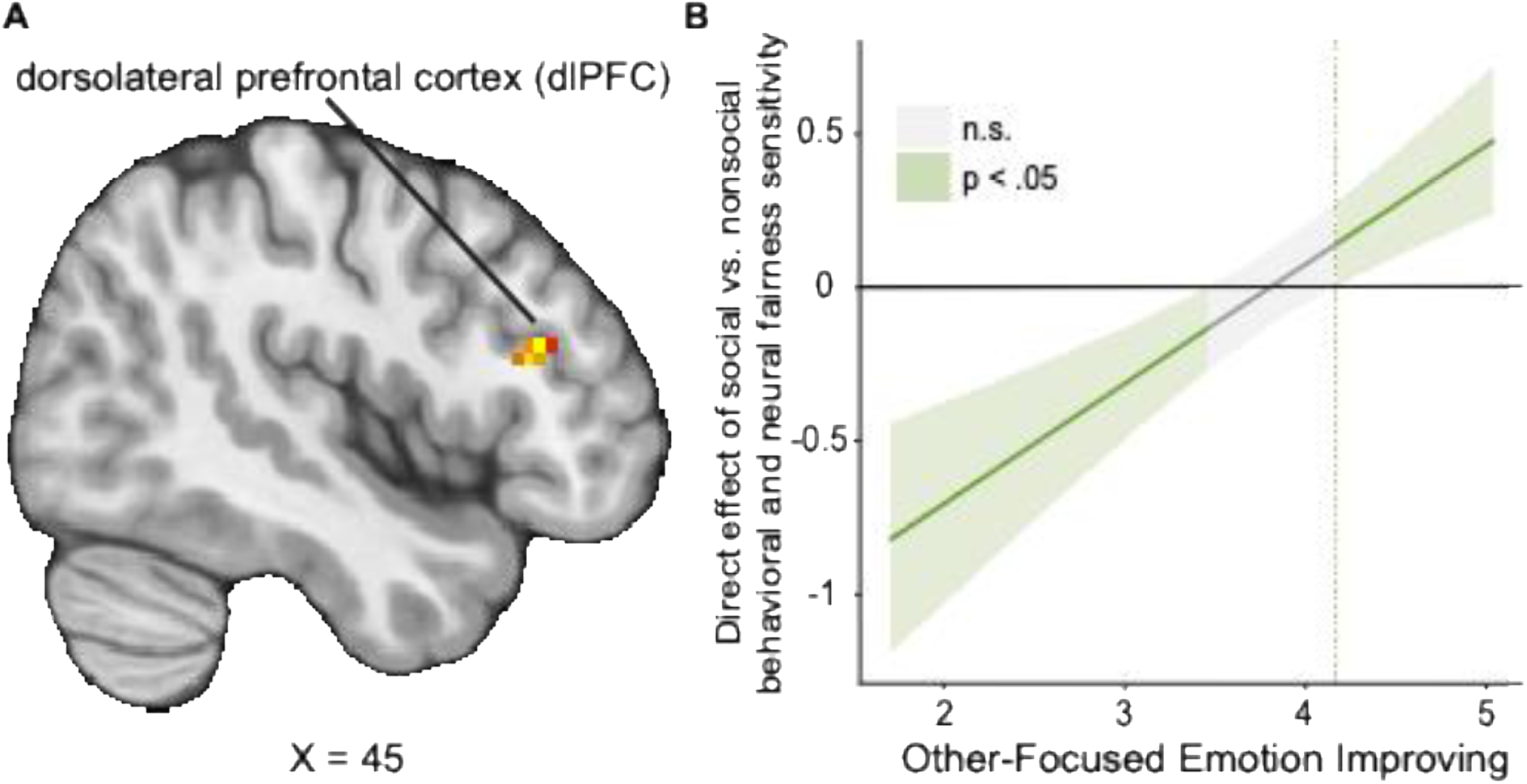
Other-directed emotion improving moderates the relationship between dlPFC activation and context-dependent behavioral sensitivity to fairness. Participants who more strongly endorse efforts to make others feel better show greater right dorsolateral prefrontal cortex (dlPFC) activation as their behavioral sensitivity to fairness in social relative to nonsocial contexts increases. That is, as individuals more frequently reported attempting to better others’ feelings, dlPFC activation was elevated when deciding to punish unfair offers from a human partner as compared to the computer partner. Further, individuals lower in other-directed emotion improving abilities also had heightened activation in the region when rejecting unfair offers from the computer as compared to the stranger. Panel A depicts the extracted dlPFC from a whole-brain contrast of social context and parametrically-modulated offer fairness with other-directed emotion improving as a predictor (Thresholded: https://identifiers.org/neurovault.image:1028050, Unthresholded: https://identifiers.org/neurovault.image:1028049). Panel B shows a Johnson-Neyman plot of this effect; shaded regions represent 95% confidence intervals with non-significant regions of the regression line in gray.

### Emotion regulation changes how affective and social information are integrated in decision-making

To examine how emotion regulation influences neural coordination among regions responsive to fairness and social context, we focused on task-dependent connectivity among key affective and social systems. For seed regions, we used the left anterior insula and bilateral amygdala as identified in our above whole-brain activation analyses of social > nonsocial trials (amygdala) and the effect of unfairness (anterior insula; see Supplemental).

Given the amygdala’s strong response to social context in our activation analyses, we next used it as a seed region in parallel PPI models. We first tested whether fairness-related amygdala connectivity differed between human and computer partners at the group-level and did not observe any significant clusters at our correction threshold. We then examined whether individual differences in socially context-dependent behavioral fairness sensitivity were associated with fairness-related amygdala connectivity. Here, individuals with higher social vs. nonsocial behavioral sensitivity to fairness—those who were more likely to reject unfair offers from human as compared to computer partners—showed stronger amygdala–orbitofrontal coupling (OFC; *z* = 3.77, *k* = 16; *p* = .04; Figure 4). This increased amygdala-OFC connectivity suggests that individuals more sensitive to fairness in social contexts may more strongly integrate social- and value-based information into their decisions to punish norm violations.

**Figure 4.**
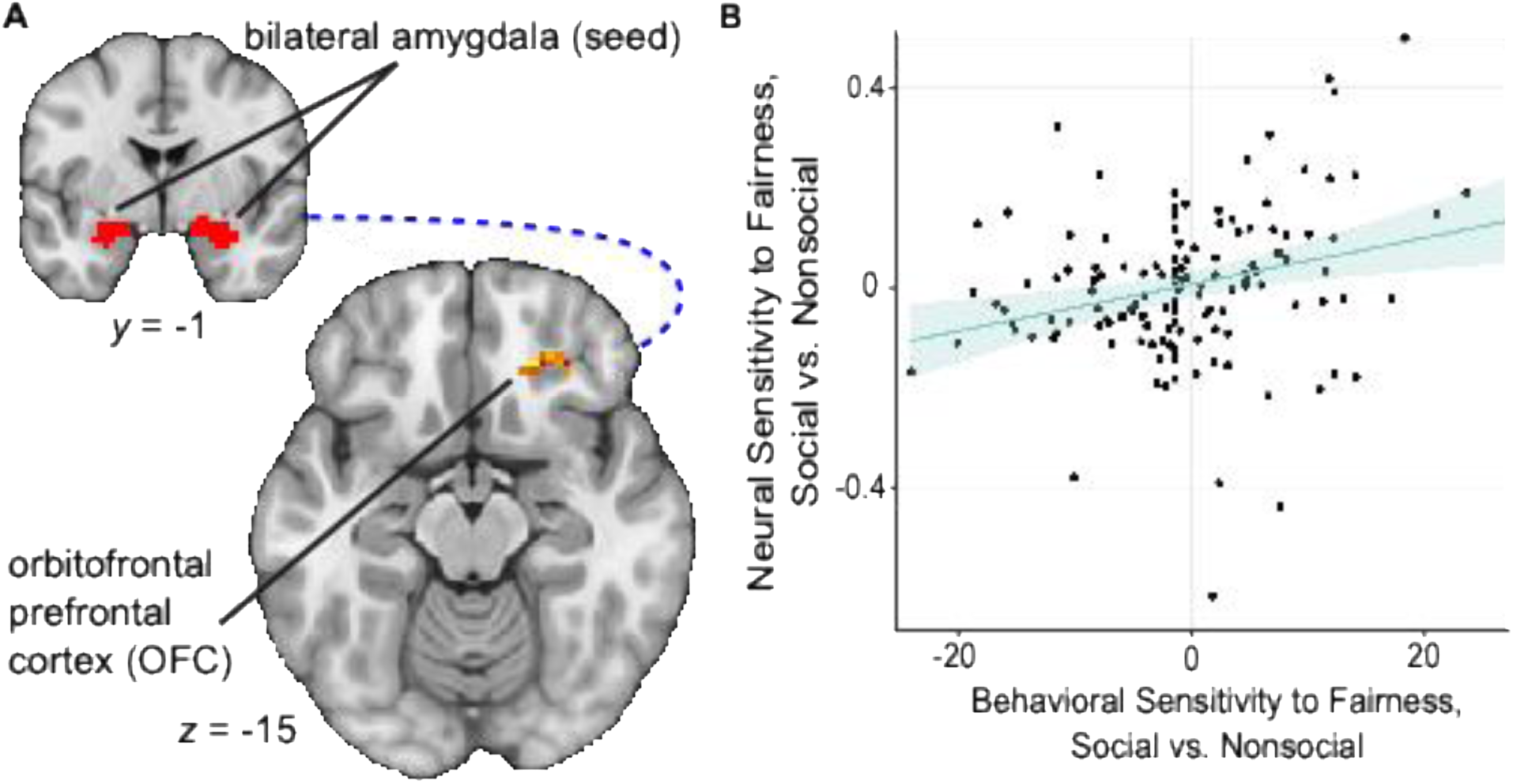
Amygdala-OFC connectivity is associated with context-dependent behavioral sensitivity to fairness. Connectivity between the bilateral amygdala seed and target orbitofrontal cortex (OFC) was positively associated with individuals’ social vs. nonsocial behavioral sensitivity to fairness. That is, as participants more frequently rejected unfair offers from human as compared to computer partners, coupling between the amygdala-OFC increased. These results suggest that amygdala-OFC connectivity is associated with individuals’ decisions to punish social partners for unfair offers. Panel A depicts the seed amygdala (inset) and OFC extracted from a whole-brain contrast of social context and parametrically-modulated offer fairness (Thresholded: https://identifiers.org/neurovault.image:1028052, Unthresholded: https://identifiers.org/neurovault.image:1028051). Panel B depicts the partner-dependent association between neural and behavioral sensitivity to fairness; shaded regions are 95% CIs.

Building on this association between social behavioral fairness sensitivity and fairness-related amygdala connectivity, we also tested whether emotion regulation tendencies further were associated with amygdala coupling during unfair social decisions. In this moderation analysis, we examined whether the link between context-dependent behavioral sensitivity to fairness and fairness-related amygdala connectivity varied with participants’ self- or other-directed emotion regulation tendencies. Among individuals with higher behavioral sensitivity to fairness in the social relative to nonsocial condition, those who more often worsened others’ emotions showed stronger fairness-related coupling between the amygdala and dorsomedial prefrontal cortex (dmPFC; *k* = 26, *p* = .003, adjusted *p* = .012; Figure 5). This pattern indicates that fairness-related amygdala–dmPFC connectivity varies systematically with both behavioral sensitivity to fairness and other-directed emotion worsening, explicitly linking interpersonal emotion regulation to neural regions previously implicated in social and self-directed emotion regulation processes. There were no comparable moderation effects observed for any of the other three emotion regulation tendencies assessed following FWER correction.

**Figure 5.**
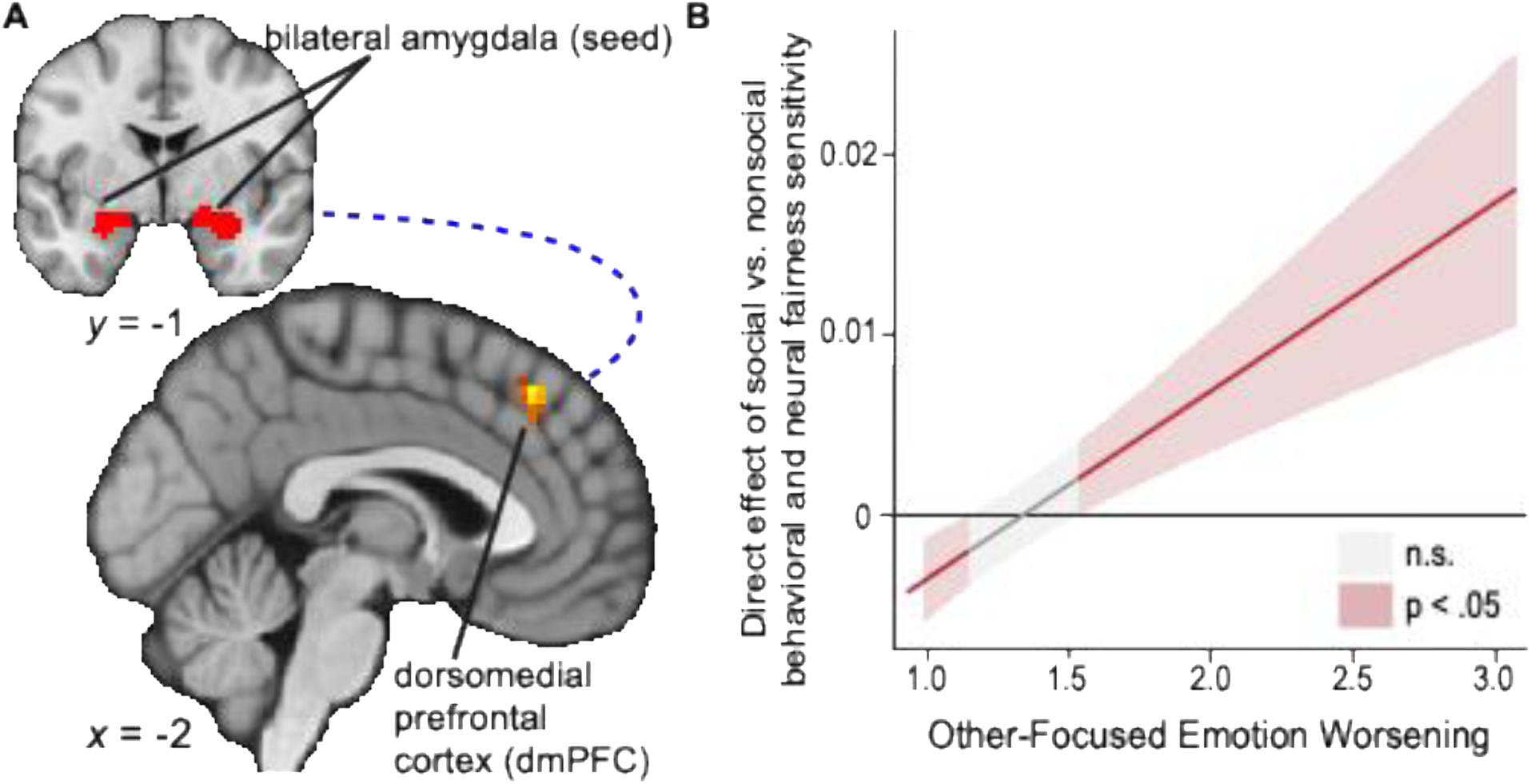
Other-directed affect worsening moderates the relationship between amygdala-dmPFC connectivity and context-dependent behavioral sensitivity to fairness. Participants who more strongly endorse efforts to make others feel worse show greater connectivity between the bilateral amygdala and dorsomedial prefrontal cortex (dmPFC) as they increasingly make decisions to punish unfair social vs. nonsocial partners. That is, as individuals more frequently reported attempting to worsen others’ feelings, amygdala-dmPFC connectivity was elevated when deciding to punish unfair offers from a human partner as compared to the computer partner. Panel A depicts the seed amygdala (inset) and dmPFC extracted from a whole-brain contrast of social context and parametrically-modulated offer fairness with other-directed emotion worsening as a predictor (Thresholded: https://identifiers.org/neurovault.image:1028055, Unthresholded: https://identifiers.org/neurovault.image:1028053). Panel B depicts the partner-dependent association between neural and behavioral sensitivity to fairness; shaded regions are 95% CIs.

With the anterior insula as a seed region, as with the amygdala, we first tested whether its connectivity with the rest of the brain varied with offer fairness and whether these fairness-related changes differed between social and nonsocial partners. At our cluster-corrected threshold of *Z* > 3.1, this analysis did not reveal any regions whose fairness-related connectivity with anterior insula differed between partner conditions. We then tested whether anterior insula connectivity tracked individual differences in behavioral sensitivity to fairness or emotion regulation tendencies and again found no significant associations or moderation effects for the anterior insula seed. These null results suggest that anterior insula connectivity during fairness-related processes may operate similarly across individuals regardless of social context or emotion regulation tendencies.

## Discussion

Perceived fairness shapes how we evaluate others and decide whether to accept or punish them. We extend prior UG work by showing that other-directed emotion regulation tendencies relate to how strongly fairness predicts choice, with individuals who prefer to worsen others’ emotions demonstrating greater behavioral sensitivity to unfair offers. Neural analyses revealed that unfairness engaged both affective (e.g., amygdala, anterior insula) and executive function (e.g., dlPFC) circuitry in trials in which participants believed that they were playing with a human partner. This was the case even though behavioral acceptance was similar across social (human) vs. nonsocial (computer) conditions. Moreover, as participants in the social vs. nonsocial condition more often rejected unfair offers, individual differences in other-directed emotion improving moderated heightened activation in the dlPFC. In the same condition, heightened connectivity between the amygdala and OFC was associated with increased rejection of unfair offers. Individual differences in other-directed emotion worsening moderated connectivity between the amygdala and dmPFC. Taken together, these findings suggest that although behavior appeared uniform across contexts, social context and emotion regulation tendencies are potentially neurally encoded in circuitry supporting valuation, norm processing, and social cognition.

A well-established behavioral signature of the UG is that people frequently reject unfair offers, even at a cost to themselves. This is often interpreted as individuals demonstrating sensitivity to fairness and enforcing socially-established norms (Gabay et al., 2014). Our finding that anterior insula activity increased with unfairness more strongly in social as compared to nonsocial contexts aligns with the insula’s proposed multifaceted role as both an “inequality detector” and salience encoder (Tabibnia et al., 2008; Molnar-Szakacs & Uddin, 2022; Huang et al., 2021). This is consistent with prior work finding that the anterior insula signals inequality and deviations from fairness norms (Cheng et al., 2017; Yang et al., 2022) and lesion evidence suggests a causal role for the insula in aversion to social inequity (Nitsch et al., 2021). Together, these findings are consistent with accounts linking anterior insula to unfairness and salience of norm violations. Our findings extend this pattern and indicate that social context modulates neural responses to unfairness even when behavioral punishment rates are similar across partners.

Among individuals more inclined to improve others’ emotions, right dlPFC activation was greater when deciding to punish unfair offers from human relative to computer partners, suggesting that the interpersonal context of the decision modulated recruitment of the region. This pattern is consistent with the dlPFC’s noted role in goal-directed control over social behavior, particularly when a trait-like tendency to better others’ emotions comes into conflict with fairness-based social norm enforcement. The dlPFC has been shown to track inequality signals in ways consistent with computational models of social preference, suggesting that it encodes deviations from expected fairness that inform subsequent accept-reject decisions (Holper et al., 2018). Converging causal evidence from noninvasive brain stimulation further supports a role for the right dlPFC in implementing norm-consistent behavior, as increasing right dlPFC excitability increases willingness to punish unfair behavior (Ruff et al., 2013). Beyond norm enforcement, dlPFC recruitment also scales during social decisions that require integration of multiple decision variables, including fairness norms, potential payoffs, and expectations about others’ behavior (Sazhin et al., 2024). The dlPFC thus may be integrating these factors to guide strategic social choice.

Individuals who were more sensitive to fairness in social relative to nonsocial contexts showed stronger amygdala-OFC connectivity during their decisions to reject unfair offers from human partners, and among those individuals, an increased tendency to worsen others’ emotions was associated with greater amygdala-dmPFC coupling. Together, these connectivity patterns suggest that context-dependent sensitivity to fairness and other-direction emotion regulation tendencies are both reflected in how the amygdala coordinates with prefrontal regions during decisions to punish unfairness. The amygdala is prototypically involved in emotion generation and rapid rejection responses to unfairness; Gospic et al. (2011) showed that amygdala activation contributes to the immediate rejection of unfair offers in the UG, implicating a region prototypically associated with affect in punishment decisions. More broadly, the amygdala integrates contextual and affective information relevant to social evaluation (Šimić et al., 2021; FeldmanHall et al., 2019), and primate evidence suggests that amygdala neurons encode socially relevant decision variables during interactive choice selection (S. W. C. Chang et al., 2015).

One possibility is that the amygdala-OFC and amygdala-dmPFC connectivity findings track distinguishable processes: one oriented toward punishing the unfair offer itself to correct a social norm violation, and one oriented towards evaluating and punishing the specific social agent responsible for it. The amygdala-OFC finding is consistent with accounts of the OFC as supporting a goal-directed cognitive map that integrates social context to guide behavior (Shi et al., 2023). In contrast, the amygdala-dmPFC connectivity observed may be more reflective of how individuals evaluate the specific social partner. The dmPFC has been implicated in impression formation and encoding person-specific information during social evaluation to support nuanced representations of others’ intentions (Ferrari et al., 2016; Grossmann & Allison, 2024; Reddan et al., 2025), and within social decision-making, its engagement has been linked to integrating social cues about others’ motives and reputation (Clairis & Lopez-Persem, 2023). This moderation by other-directed emotion worsening may indicate that unfair offers from human partners are sometimes processed not just as violations of socially-established fairness norms to be corrected, as with the OFC connectivity finding, but are also related to the evaluation of the specific social agent who committed the transgression.

These findings may also be viewed through moral psychology accounts that distinguish evaluating unfair outcomes from evaluating unfair agents. People are not only sensitive to the fact that a social norm has been violated, but also to who committed the violation and why, a distinction that can shape both affective responses and subsequent punitive decisions (Helion et al., 2020; Helion & Ochsner, 2018). In this framework, fairness-related responses may reflect both sensitivity to the unfairness of an outcome and evaluations of the intentions or character of the individual responsible for the violation. In line with this account, our observation that other-directed emotion worsening moderated amygdala-dmPFC connectivity in response to unfair social offers suggests that some individuals may be especially attuned to evaluating the agent committing the perceived fairness violation and not just the violation itself. Together, these findings suggest that individual differences in emotion regulation tendencies in fairness- related decisions shape the neural encoding of fairness violations and social evaluations associated with the interaction.

Despite these advances, several caveats warrant consideration. First, although our use of one-shot trials minimized strategic learning and reputational concerns (L. J. Chang & Sanfey, 2009), it may have reduced participants’ motivation to reject unfair offers. Indeed, work using multi-period and iterated ultimatum games has shown that expectations, offers, and rejection behavior can adapt over time as players respond to one another’s choices (Azar et al., 2015; Shaw et al., 2018). Second, because we did not collect trial-by-trial ratings of how participants perceived each partner, we were unable to assess whether factors like closeness or social affinity impacted decision-making. Future studies could incorporate partner-perception measures or log responses to partner images to address this gap. Finally, our emotion-regulation measure captured self-reported tendencies (Niven et al., 2011), which align with aspects of participants’ behavior but do not necessarily index moment-to-moment regulation during decision-making (Jung et al., 2025). Tasks that explicitly instruct participants to regulate others’ as well as their own emotions may help identify the processes through which emotion regulation shapes fairness-related decisions in social contexts.

In conclusion, the present study identified other-directed emotion regulation abilities, as opposed to self-directed regulation, as a meaningful influence on social decision-making. While prior work has largely looked at instructed, self-focused reappraisal paradigms, the current findings demonstrate that trait-like tendencies to influence others’ emotions are associated with both behavioral sensitivity to fairness and neural responsivity and connectivity that supports punishment decisions. Importantly, though behavior did not significantly vary between social and nonsocial contexts, neural analyses indicated that social context was still encoded. This supports the idea that fairness-related decisions are embedded within socially-sensitive affective and cognitive systems, even when observed behavioral decisions appear uniform. Together, these findings suggest that other-directed emotion regulation may help to explain reliable individual differences in fairness-related behavior and neural responsivity. These results motivate future work examining how partner perceptions and repeated interactions shape the link between interpersonal emotion regulation and fairness in social decision-making contexts.

## Acknowledgments

The authors thank Daniel Sazhin, Cooper J. Sharp, and Abraham Dachs for assistance with data collection and data analysis.

## Data and code availability

Analysis code can be found on GitHub (https://github.com/DVS-Lab/ugr-emoreg). All statistical maps (thresholded and unthresholded) and masks used for analyses can be found on NeuroVault (https://identifiers.org/neurovault.collection:23412); and preliminary raw neuroimaging data can be found on OpenNeuro (doi:10.18112/openneuro.ds005123.v1.1.2).

