## Supplemental Information for "Differences in other-directed emotion regulation tracks connectivity between amygdala and prefrontal regions during fairness decisions"

#### Methods

##### Behavioral Analyses

To ensure participants only viewed offers in whole dollar amounts, the 5% and 10% offer amounts were slightly adjusted. For the low endowment condition ($16), these corresponded to $1 and $2, respectively. For the high endowment condition ($32), these corresponded to $2 and $3, respectively.

To estimate behavioral sensitivity to fairness, offer fairness was input at the trial level, mean-centered, and treated as a continuous variable. Participant-specific slopes for behavioral sensitivity to fairness were calculated with a generalized linear mixed effects model (GLMM) with a binomial outcome variable (accept or reject), fixed and random slopes for offer fairness, and a random intercept for participant. Participant-specific slopes were computed as the sum of the fixed effect and participant-level random effect for offer fairness. Models were estimated separately for 1) social condition trials only (i.e., stranger as partner), 2) nonsocial condition trials only (i.e., computer as partner), and 3) across all trials (i.e., collapsed across both social and nonsocial partner conditions). The following formula was utilized on trial-level data:

$$glmer(accept \sim behavioral\ fairness + (1 +\ behavioral\ fairness \mid subject)$$

where “accept” are binary accept/reject decisions, behavioral fairness is the percentage of the endowment offered (i.e., 5%, 10%, 25%, or 50%), and subject is the participant. Slope values are the sum of the fixed effect (i.e., behavioral fairness) and random effect (i.e., behavioral fairness with random intercept for participant), in line with best linear unbiased prediction. Slopes for social trials only, nonsocial trials only, and all trials were calculated for all subjects, as well as social vs. nonsocial difference slope composite value by subtracting each subject's nonsocial slope from their social slope.

#### Neural Analyses

PPI seeds for amygdala and anterior insula (AIns) were extracted from activation-based results. Specifically, the bilateral amygdala seed was defined as the overlapping regions from the Harvard-Oxford Atlas and results from the social vs. nonsocial activation contrast. The AIns seed was defined from the parametrically modulated fairness activation contrast and extracted using a 5mm sphere using MRIcron (Rorden, 2025). The AIns was selected as it was the most significant cluster associated with offer unfairness-related responses, as described above, whereas the amygdala was selected as it was the most significant cluster during human vs. computer trials, save for the fusiform face area. Cluster results can be found in Tables 2 and 3.

Activation and PPI-based analyses were corrected for multiple comparisons in FSL. All analyses looked at neural responses during the decision phase in the social vs. nonsocial condition with parametrically modulated offer fairness. Cluster results can be found in Table 4.

### Imaging pre-processing.

#### Creation of B0 Field maps (warpkit)

We created static B0 field maps from the phase information in our multi-echo functional MRI (ME-fMRI) echo-planar imaging (EPI) data using the Multi-Echo DIstortion Correction (MEDIC) algorithm (version 0.1.1; https://github.com/vanandrew/warpkit) (Van et al., 2023). This method estimates voxelwise B0 inhomogeneity by modeling the phase of the MRI signal as a linear function of echo time (Jezzard & Balaban, 1995). To remove channel-specific phase offsets introduced during multi-coil image reconstruction, we applied Multi-Channel Phase Combination using 3D Simultaneous estimation (MCPC-3D-S), which derives a zero-time phase offset based on the unwrapped phase difference between the first two echoes (Eckstein et al., 2018). Phase unwrapping was then performed using Rapid Open-source Minimum-spanning-tree phase unwrapping (ROMEO), which jointly unwraps all echoes under the constraint that phase accumulates linearly with echo time, thereby improving stability and accuracy (Dymerska et al., 2021). The resulting unwrapped phase data were fitted using a magnitude-weighted least-squares approach to compute off-resonance field maps in units of hertz, which were then transformed to anatomical space for use in susceptibility distortion correction.

#### fMRIPrep

Results included in this manuscript come from preprocessing performed using *fMRIPrep* 24.1.1 (Esteban et al. (2019); Esteban et al. (2018); RRID:SCR_016216), which is based on *Nipype* 1.8.6 (K. Gorgolewski et al. (2011); K. J. Gorgolewski et al. (2018); RRID:SCR_002502).

Preprocessing of $B_0$ inhomogeneity mappings

A total of 8 fieldmaps were found available within the input BIDS structure for this particular subject. A $B_0$ nonuniformity map (or *fieldmap*) was directly measured with an MRI scheme designed with that purpose such as SEI (Spiral-Echo Imaging).

<u>Anatomical data preprocessing</u>

A total of 1 T1-weighted (T1w) images were found within the input BIDS dataset. The T1w image was corrected for intensity non-uniformity (INU) with N4BiasFieldCorrection (Tustison et al. 2010), distributed with ANTs 2.5.3 (Avants et al. 2008, RRID:SCR_004757), and used as T1w-reference throughout the workflow. The T1w-reference was then skull-stripped with a *Nipype* implementation of the antsBrainExtraction.sh workflow (from ANTs), using OASIS30ANTs as target template. Brain tissue segmentation of cerebrospinal fluid (CSF), white-matter (WM) and gray-matter (GM) was performed on the brain-extracted T1w using fast (FSL (version unknown), RRID:SCR_002823, Zhang, Brady, and Smith 2001). Volume-based spatial normalization to two standard spaces (MNI152NLin6Asym, MNI152NLin2009cAsym) was performed through nonlinear registration with antsRegistration (ANTs 2.5.3), using brain-extracted versions of both T1w reference and the T1w template. The following templates were were selected for spatial normalization and accessed with *TemplateFlow* (24.2.0, Ciric et al. 2022): *FSL's MNI ICBM 152 non-linear 6th Generation Asymmetric Average Brain Stereotaxic Registration Model* [Evans et al. (2012), RRID:SCR_002823; TemplateFlow ID: MNI152NLin6Asym], *ICBM 152 Nonlinear Asymmetrical template version 2009c* [Fonov et al. (2009), RRID:SCR_008796; TemplateFlow ID: MNI152NLin2009cAsym].

<u>Functional data preprocessing</u>

For each of the 8 BOLD runs found per subject (across all tasks and sessions), the following preprocessing was performed. First, a reference volume was generated from the shortest echo of the BOLD run, using a custom methodology of *fMRIPrep*, for use in head motion correction. Head-motion parameters with respect to the BOLD reference (transformation matrices, and six corresponding rotation and translation parameters) are estimated before any spatiotemporal filtering using mcflirt (FSL, Jenkinson et al. 2002). The estimated *fieldmap* was then aligned with rigid-registration to the target EPI (echo-planar imaging) reference run. The field coefficients were mapped on to the reference EPI using the transform. The BOLD reference was then co-registered to the T1w reference using mri_coreg (FreeSurfer) followed by flirt (FSL , Jenkinson and Smith 2001) with the boundary-based registration (Greve and Fischl 2009) cost-function. Co-registration was configured with six degrees of freedom. Several confounding time-series were calculated based on the *preprocessed BOLD*: framewise displacement (FD), DVARS and three region-wise global signals. FD was computed using two formulations following Power (absolute sum of relative motions, Power et al. (2014)) and Jenkinson (relative root mean square displacement between affines, Jenkinson et al. (2002)). FD and DVARS are calculated for each functional run, both using their implementations in *Nipype* (following the definitions by Power et al. 2014). The three global signals are extracted within the CSF, the WM, and the whole-brain masks. Additionally, a set of physiological regressors were extracted to allow for component-based noise correction (*CompCor*, Behzadi et al. 2007). Principal components are estimated after high-pass filtering the *preprocessed BOLD* time-series (using a discrete cosine filter with 128s cut-off) for the two *CompCor* variants: temporal (tCompCor) and anatomical (aCompCor). tCompCor components are then calculated from the top 2% variable voxels within the brain mask. For aCompCor, three probabilistic masks (CSF, WM and combined CSF+WM) are generated in anatomical space. The implementation differs from that of Behzadi et al. in that instead of eroding the masks by 2 pixels on BOLD space, a mask of pixels that likely contain a volume fraction of GM is subtracted from the aCompCor masks. This mask is obtained by thresholding the corresponding partial volume map at 0.05, and it ensures components are not extracted from voxels containing a minimal fraction of GM. Finally, these masks are resampled into BOLD space and binarized by thresholding at 0.99 (as in the original implementation). Components are also calculated separately within the WM and CSF masks. For each CompCor decomposition, the *k* components with the largest singular values are retained, such that the retained components' time series are sufficient to explain 50 percent of variance across the nuisance mask (CSF, WM, combined, or temporal). The remaining components are dropped from consideration. The head-motion estimates calculated in the correction step were also placed within the corresponding confounds file. The confound time series derived from head motion estimates and global signals were expanded with the inclusion of temporal derivatives and quadratic terms for each (Satterthwaite et al. 2013). Frames that exceeded a threshold of 0.5 mm FD or 1.5 standardized DVARS were annotated as motion outliers. Additional nuisance timeseries are calculated by means of principal components analysis of the signal found within a thin band (*crown*) of voxels around the edge of the brain, as proposed by (Patriat, Reynolds, and Birn 2017). All resamplings can be performed with *a single interpolation step* by composing all the pertinent transformations (i.e. head-motion transform matrices, susceptibility distortion correction when available, and co-registrations to anatomical and output spaces). Gridded (volumetric) resamplings were performed using nitransforms, configured with cubic B-spline interpolation.

Many internal operations of *fMRIPrep* use *Nilearn* 0.10.4 (Abraham et al. 2014, RRID:SCR_001362), mostly within the functional processing workflow. For more details of the pipeline, see the section corresponding to workflows in *fMRIPrep*'s documentation.

Copyright Waiver

The above boilerplate text was automatically generated by fMRIPrep with the express intention that users should copy and paste this text into their manuscripts *unchanged*. It is released under the CC0 license.

#### Echo-Time Dependent Denoising (tedana)

TE-dependence analysis was performed on input data using the tedana workflow (DuPre et al. 2021). An initial mask was generated from the first echo using nilearn's compute_epi_mask function. An adaptive mask was then generated using the dropout method(s), in which each voxel's value reflects the number of echoes with 'good' data. A two-stage masking procedure was applied, in which a liberal mask (including voxels with good data in at least the first echo) was used for optimal combination, T2*/S0 estimation, and denoising, while a more conservative mask (restricted to voxels with good data in at least the first three echoes) was used for the component classification procedure. A monoexponential model was fit to the data at each voxel using nonlinear model fitting in order to estimate T2* and S0 maps, using T2*/S0 estimates from a log-linear fit as initial values. For each voxel, the value from the adaptive mask was used to determine which echoes would be used to estimate T2* and S0. In cases of model fit failure, T2*/S0 estimates from the log-linear fit were retained instead. Multi-echo data were then optimally combined using the T2* combination method (Posse et al. 1999). Principal component analysis based on the PCA component estimation with a Moving Average (stationary Gaussian) process (Li et al. 2007) was applied to the optimally combined data for dimensionality reduction. The following metrics were calculated: kappa, rho, countnoise, countsigFT2, countsigFS0, dice_FT2, dice_FS0, signal-noise_t, variance explained, normalized variance explained, d_table_score. Kappa (kappa) and Rho (rho) were calculated as measures of TE-dependence and TE-independence, respectively. A t-test was performed between the distributions of T2*-model F-statistics associated with clusters (i.e., signal) and non-cluster voxels (i.e., noise) to generate a t-statistic (metric signal-noise_z) and p-value (metric signal-noise_p) measuring relative association of the component to signal over noise. The number of significant voxels not from clusters was calculated for each component. Independent component analysis was then used to decompose the dimensionally reduced dataset. The following metrics were calculated: countnoise, countsigFS0, countsigFT2, d_table_score, dice_FS0, dice_FT2, kappa, normalized variance explained, rho, signal-noise_t, variance explained. Kappa (kappa) and Rho (rho) were calculated as measures of TE-dependence and TE-independence, respectively. A t-test was performed between the distributions of T2*-model F-statistics associated with clusters (i.e., signal) and non-cluster voxels (i.e., noise) to generate a t-statistic (metric signal-noise_z) and p-value (metric signal-noise_p) measuring relative association of the component to signal over noise. The number of significant voxels not from clusters was calculated for each component.

Next, component selection was performed to identify BOLD (TE-dependent) and non-BOLD (TE-independent) components using a decision tree.

This workflow used numpy (Van Der Walt et al. 2011), scipy (Virtanen et al. 2020), pandas (McKinney et al. 2010, pandas development team et al. 2020), scikit-learn (Pedregosa et al. 2011), nilearn, bokeh (Team et al. 2018), matplotlib (Hunter et al. 2007), and nibabel (Brett et al. 2019). This workflow also used the Dice similarity index (Dice et al. 1945, Sorensen et al. 1948).

#### Outlier Exclusion

Runs were determined to be outliers based on a standard boxplot threshold applied to framewise displacement (FD) and temporal signal-to-noise ratio (tSNR). Specifically, runs were excluded if FD exceeded the upper bound defined as Q3 + 1.5*IQR or if tSNR was below the lower bound defined as Q1 – 1.5*IQR.

#### Additional Figures

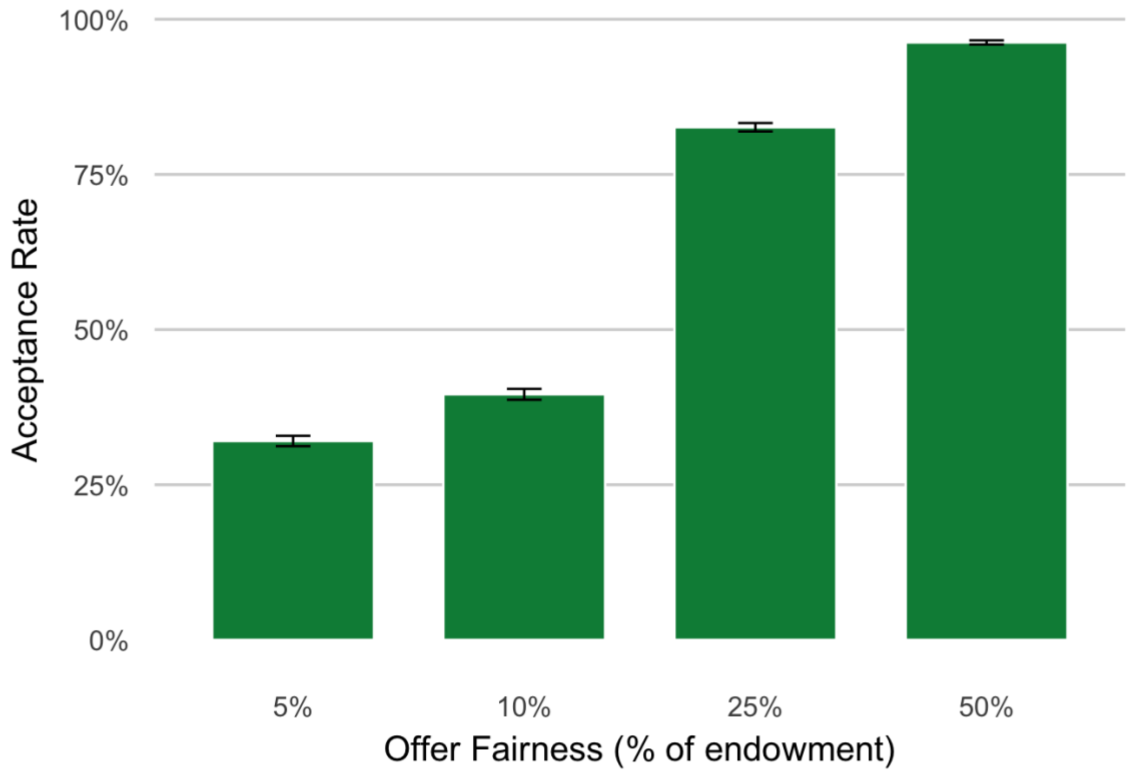


**Supplemental Figure 1.** Increased offer fairness is associated with increased acceptance rates in participants, $b = 15.80$, $SE = 0.34$, $z = 47.0$ $p < .001$. There was no main effect of partner condition nor significant interaction of offer fairness with partner condition on offer acceptance rates.

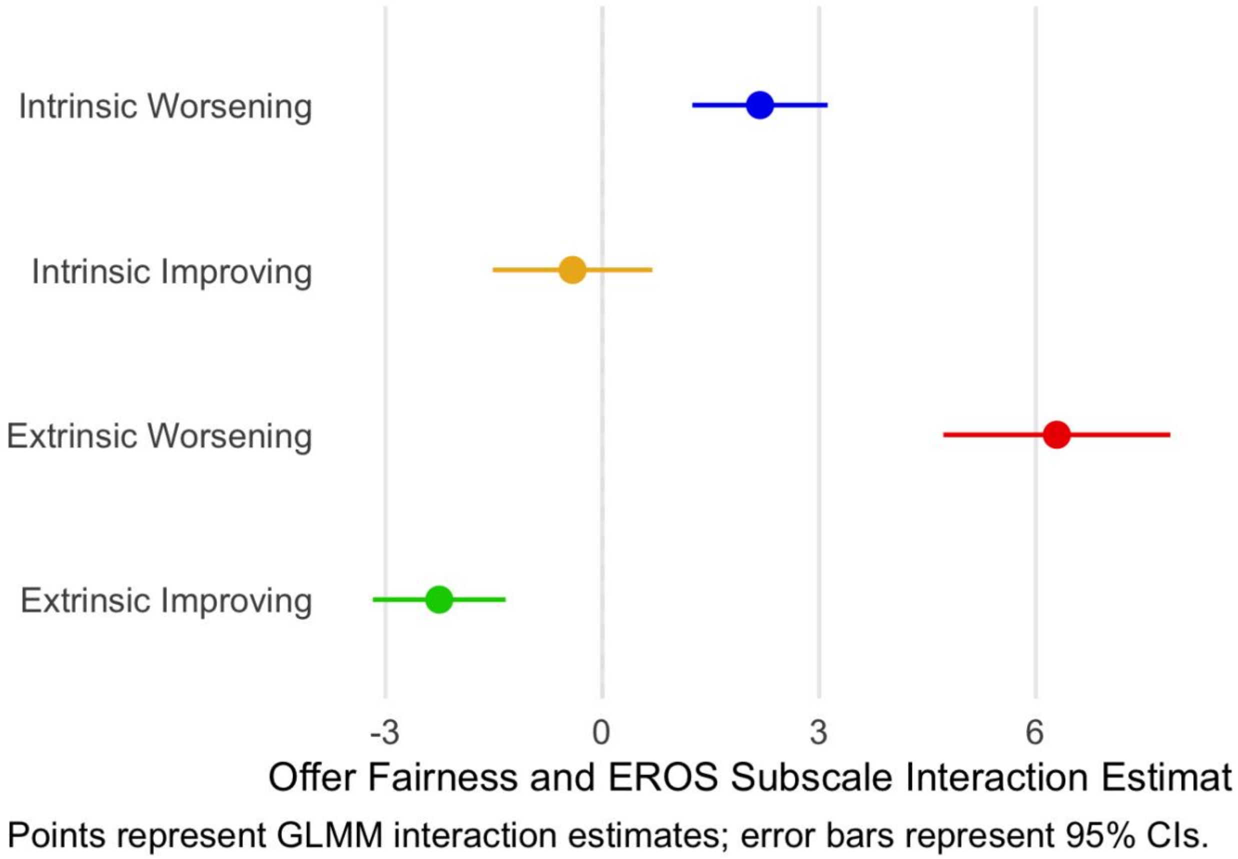


**Supplemental Figure 2.** Extrinsic affect-worsening (b = 6.29, SE = 0.80, z = 7.85, p < .001), extrinsic affect-improving (b = -2.26, SE = 0.47, z = -4.81, p < .001), and intrinsic-affect worsening (b = 2.18, SE = 0.48, z = 4.57, p < .001) significantly moderated the association between offer fairness and offer acceptance rates. Points represent GLMM interaction estimates on the log-odds scale. Individuals higher in other-directed emotion worsening are more sensitive to offer fairness and are more likely to reject unfair offers.

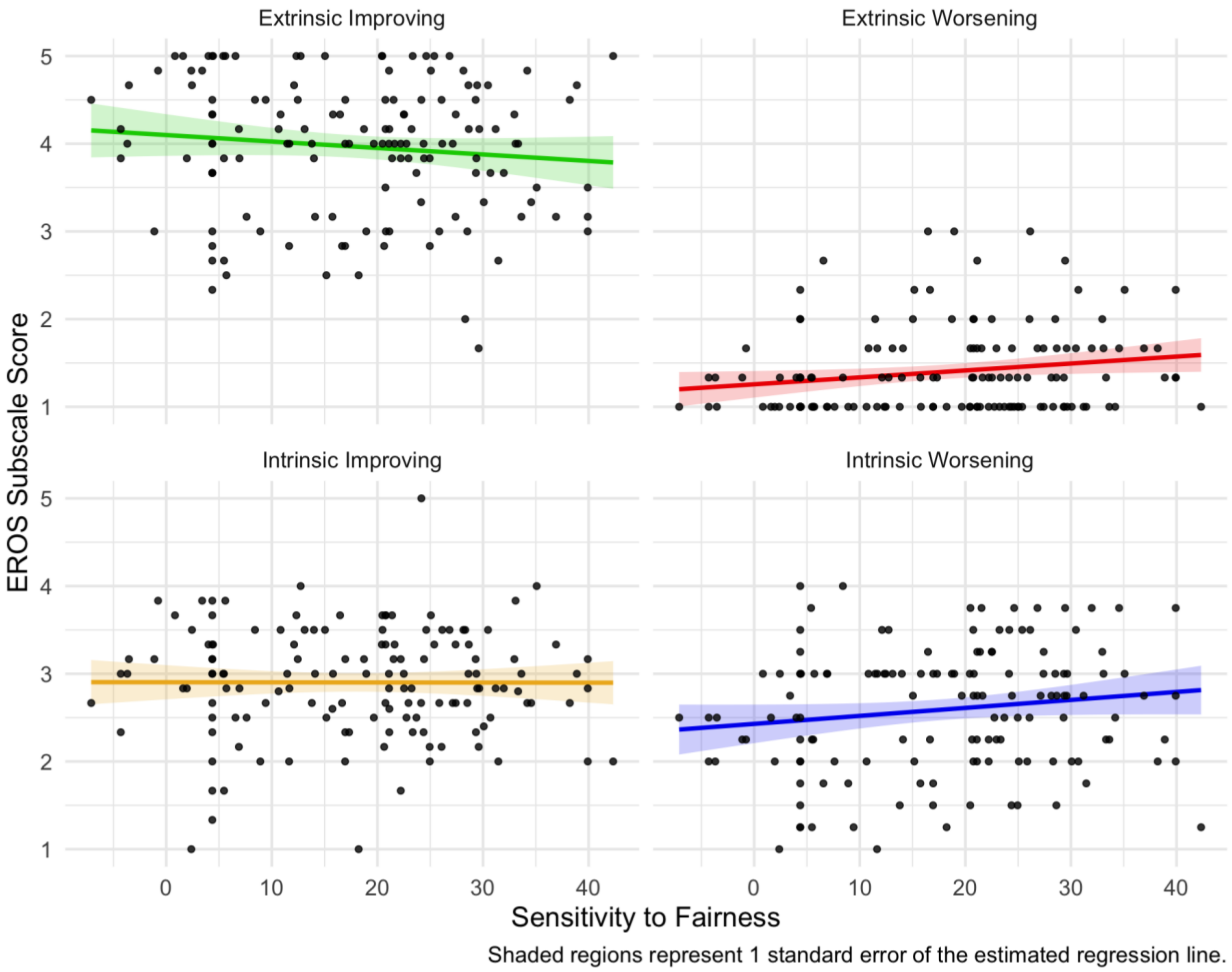


**Supplemental Figure 3.** Other-directed emotion worsening is associated with increased sensitivity to fairness, regardless of social condition, b = 4.35, SE = 1.98, t(136) = 2.20, p = .029.

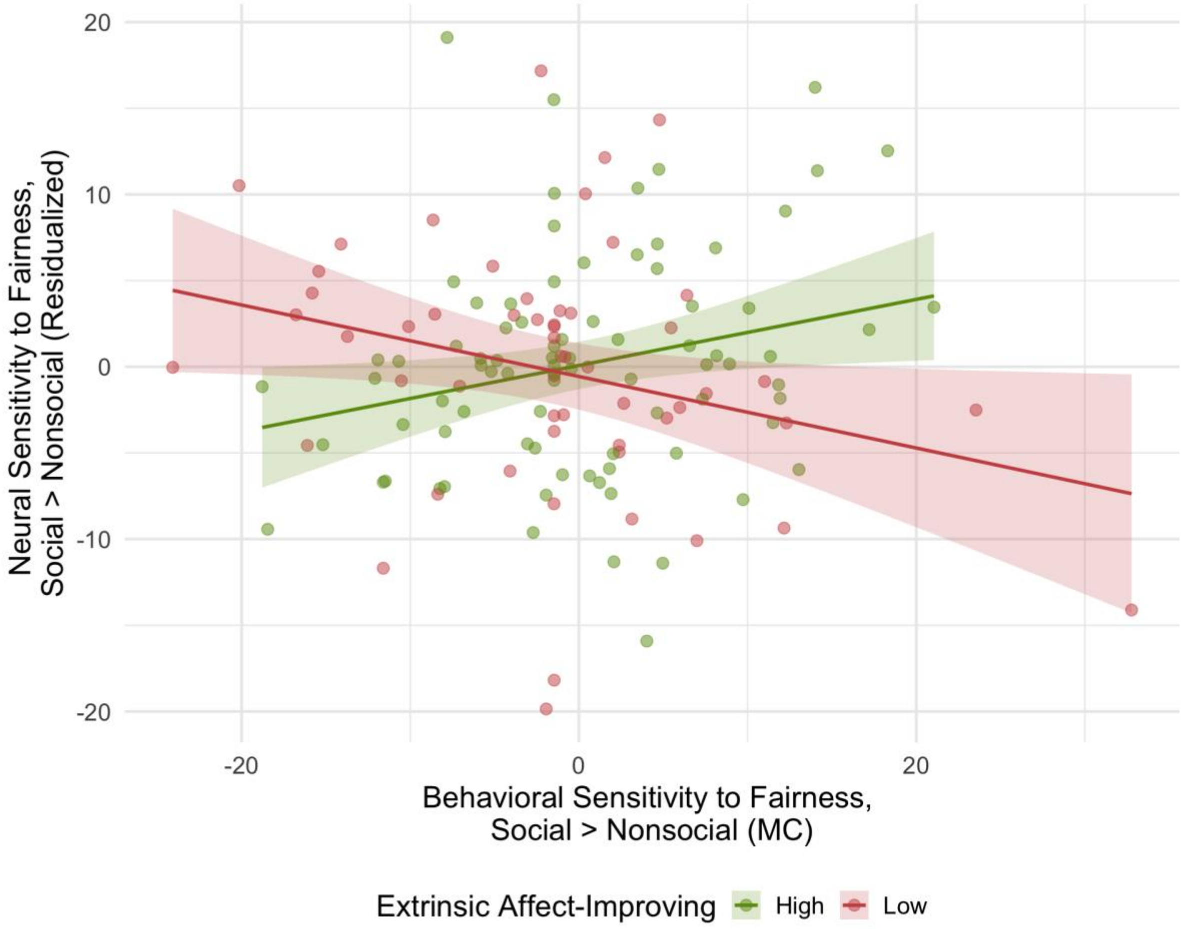


**Supplemental Figure 4**. Other-directed emotion improving moderates dlPFC activation.

Alternate presentation of the Johnson-Neyman plots used in the main manuscript.

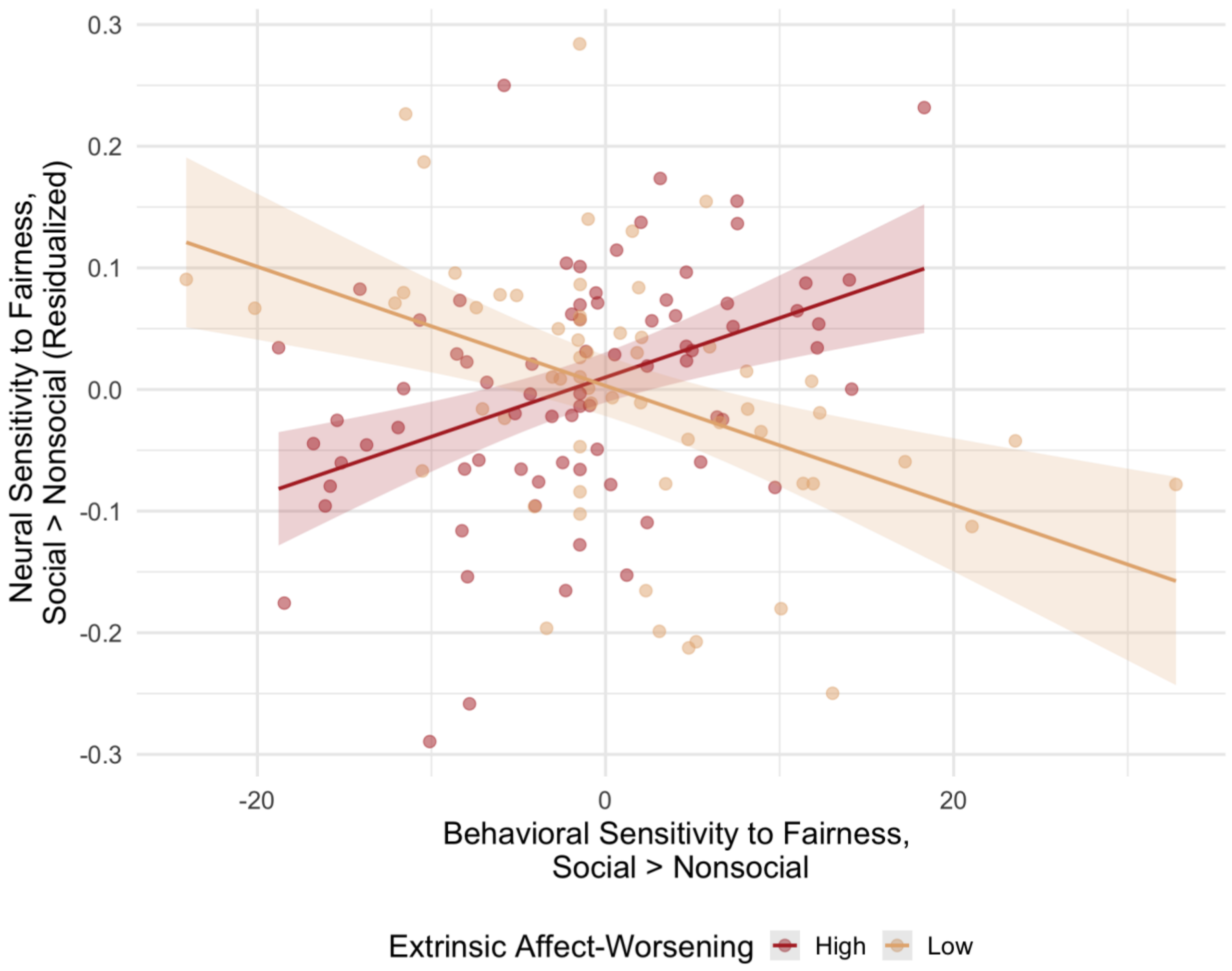


**Supplemental Figure 5.** Other-directed emotion worsening moderates amygdala-dmPFC connectivity. Alternate presentation of the Johnson-Neyman plots used in the main manuscript.

| Participant Demographics | | |
| --- | --- | --- |
| *Category* | *n* | % |
| Gender | | |
| Female | 88 | 63.8% |
| Male | 45 | 32.6% |
| Non-Binary | 5 | 3.6% |
| Race | | |
| White | 79 | 57.2% |
| Black or African American | 28 | 20.3% |
| Asian | 19 | 13.8% |
| Two or more races | 8 | 5.8% |
| Prefer not to respond | 4 | 2.9% |
| Ethnicity | | |
| Not Hispanic or Latino | 122 | 88.4% |
| Hispanic or Latino | 11 | 8% |
| Ethnicity not described (Other) | 3 | 2.2% |
| Prefer not to respond | 1 | 0.7% |

Summary reflects participants aged 21–55 in the final sample, n = 138. Gender, race, and ethnicity were self-reported.

**Table 1.** Participant demographics. Gender, race, and ethnicity information for the 138 participants aged 21-55 from the greater Philadelphia metro area used for analyses.

| Hemisphere | Region | No. of voxels | Z-max | *x* | *y* | *z* |
|---|---|---|---|---|---|---|
| **R** | FFA | 339 | 14.2 | 41.3 | -47.7 | -21.5 |
| **L** | FFA | 40 | 12.2 | -39.7 | -50.4 | -21.5 |
| **R** | Amygdala | 20 | 11.9 | 20.1 | -6.7 | -12 |
| **L** | Amygdala | 13 | 11.2 | -20.5 | -5.3 | -14.2 |
| **R** | HC | 3 | 11.2 | 16.4 | -31.9 | -1 |
| **R** | vmPFC | 4 | 10.5 | 0.2 | 47.8 | -15.7 |

**Table 2.** Social vs. Nonsocial contrast results. Voxel-height thresholding analysis with a null model (no covariates) was utilized to derive the bilateral amygdala mask. Overlap between the derived mask and anatomical Harvard-Oxford atlas mask for bilateral amygdala was used for PPI analyses. Mask was created using the positive contrast for neural response to the social vs. nonsocial condition. Table only reports results for Z-values > 10.4. Binarized mask: https://identifiers.org/neurovault.image:1023931, Full unthresholded results used to derive mask: https://identifiers.org/neurovault.image:1028143.

| Hemisphere | Region | No. of voxels | Z-max | *x* | *y* | *z* |
|---|---|---|---|---|---|---|
| **L** | aINS | 13 | 6.15 | -31.6 | 22.5 | -12.6 |
| **R** | MTG | 15 | 6.15 | 49.4 | 3.6 | -39.3 |
| **L** | mPFC | 14 | 5.95 | -4.6 | 52.2 | 32 |

**Table 3.** Offer fairness, parametrically modulated contrast results. Voxel-height thresholding analysis with a null model (no covariates) was utilized to derive the left anterior insula mask used for PPI analyses. Mask was created using the negative contrast for neural response to unfair offers. Binarized mask: https://identifiers.org/neurovault.image:1023930, Full unthresholded results used to derive mask: https://identifiers.org/neurovault.image:1028055.

| Hemisphere | Region | No. of voxels | Z-max | *x* | *y* | *z* | *p* | FWER *p* |
|---|---|---|---|---|---|---|---|---|
| | | | | **Activation** | | | | |
| **R** | dlPFC | 38 | 4.4 | 46.7 | 30.6 | 20.1 | .00134 | .00532 |
| | | | | **PPI (amygdala as seed)** | | | | |
| **L** | OFC | 16 | 3.77 | 19.7 | 36 | -15.6 | .0468 | - |
| **L** | dmPFC | 26 | 4.84 | -1.9 | 33.3 | 46.8 | .00305 | .0122 |

**Table 4.** Significant clusters from activation and PPI-based analyses in the social vs. nonsocial condition, offer fairness parametrically modulated. The dlPFC finding is moderated by other-directed emotion bettering and the dmPFC finding is moderated by other-direction emotion worsening, thus family-wise error corrected p-values are reported to account for the four potential emotion regulation tendencies as indexed by EROS.
